# Temporal Transcriptomics Identifies Isoform-specific Trans-regulation by Multiple lncRNAs in Human iPSCs

**DOI:** 10.64898/2026.05.13.724994

**Authors:** Mingfeng Liu, Izabela Mamede, Sajad Sofi, Isabela Pereira, Vishantie Dostal, Alison R. S. Pashos, Conor McMahon, Aneesh Waikar, George Stephenson, Thomas R. Cech, John L. Rinn

## Abstract

Some long non-coding RNAs (lncRNAs) are known to regulate gene expression. However, the underlying temporal dynamics of lncRNAs influencing gene and epigenetic regulation and mechanisms of lncRNA regulation in trans are less understood. To investigate this, we genetically engineered 17 doxycycline-inducible lncRNA transgenes for ectopic expression at the H11 safe harbor locus in human pluripotent stem cells (hiPSCs), and we generated high-density temporal RNA-seq and ATAC-seq profiles. Most lncRNA transgenes were induced at 2 hours and maintained expression through the 96-hour time course. Surprisingly, when we sought to identify gene expression changes due to the lncRNAs, we found that the global transcriptional landscape was dominated by a strong systemic response triggered by doxycycline exposure. We rigorously defined this cohort of genes as a Doxycycline-Responsive Gene Signature (DRGS). The DRGS was also present in at least 28 public datasets from dox-inducible transgene studies involving diverse cell types. Next, we determined which lncRNAs exhibited trans-regulatory events. We identified DANCR, FENDRR, LINC00667, LINC00847, LNCPRESS1, and PNKY as lncRNAs that regulate specific transcript expression in trans. The downstream target genes encoded 53 mRNAs and 10 lncRNAs. None of the target lncRNAs altered gene expression proximal to their own loci (i.e., triggering secondary cis-effects). Surprisingly, the target genes of LINC00847 (transcribed from chromosome 22) were substantially enriched on chromosome 19, with a preponderance of target genes encoding RNA metabolism and RNA splicing factors. Collectively, our study provides a resource to discern artifacts in the doxycycline-inducible system and identifies temporally regulated targets of 6 lncRNAs for future mechanistic studies.

## Introduction

In the past decade, hundreds of long non-coding RNAs (lncRNAs) have been characterized, leading to a multitude of mechanistic models and expanding our understanding of epigenetic gene regulation (Andergassen and Rinn 2022). These different mechanisms include lncRNA regulation of gene expression in cis and in trans, with the former regulating local chromatin structure at their transcription sites and the latter acting as mobile scaffolds or baits to regulate distant genomic sites (Huarte et al. 2010; Orom et al. 2010; Guttman and Rinn 2012; Rinn and Chang 2012; Quinodoz and Guttman 2014; Thomas and Joan 2014; Chu et al. 2015; James et al. 2015; Cheng et al. 2016; Engreitz et al. 2016; Xue et al. 2016; Chen et al. 2017; Kopp and Mendell 2018; Lewandowski et al. 2019; Consortium et al. 2020; Fang et al. 2020; Statello et al. 2021; Mattick et al. 2023). These lncRNA-mediated regulatory events are required in vivo for physiological roles ranging from fertility to development and disease (Marahrens et al. 1997; Sauvageau et al. 2013; Andersen et al. 2019; Kopp et al. 2019; Ritter et al. 2019; Rom et al. 2019; Lewandowski et al. 2020; Allou et al. 2021; Andergassen and Rinn 2022). However, these models evaluate lncRNA biology at homeostasis and therefore do not reveal the stepwise regulatory events. Whether using a gene knockout model or a stable overexpression system, phenotypic readings are typically measured days or weeks after the perturbation. This limitation makes it challenging to distinguish primary regulatory events that are directly mediated by lncRNAs from secondary or compensatory responses. To address this issue, inducible transgenic systems provide a valuable experimental approach(Gossen and Bujard 1992; Gossen et al. 1995). The temporal control in these systems reveals the order of regulatory events and the mechanistic target sites of lncRNAs in cis and trans.

The utility of such temporal resolution is exemplified in studies of lncRNA Firre in mouse embryonic stem cells. Using a time-resolved system, we showed that Firre triggers rapid de novo chromatin opening and transcription of target genes in trans within half an hour (Much et al. 2024). Moreover, Firre-mediated gene expression changes are epigenetically remembered for days. In a Firre knock-out cell line, when Firre transgene expression is induced, the aberrant transcriptomic landscape is rescued; in addition, analyses with high temporal resolution provided intermediate regulatory events that cascade to homeostasis. Because this study was performed for only one lncRNA, it raises the question whether the regulatory features observed for Firre represent a unique case or a general principle.

Here, we aimed to determine the diversity of temporal regulation of gene expression by lncRNAs in trans in human pluripotency. To this end, we developed a genetically defined hiPSC platform. We introduced a third copy of the lncRNA gene using the H11 safe harbor locus (Zhu et al. 2014) and controlled by the Tet-On system (Gossen et al. 1995) to ensure precise temporal induction. We generated 17 independent lncRNA-expressing cell lines and performed RNA-seq covering a dense time course (0 to 96 h, 8 time points) to capture gene expression trajectories and ATAC-seq(Buenrostro et al. 2013), focusing on the immediate-early phase (0 to 2.5 h, 6 time points) to understand chromatin accessibility dynamics.

Unexpectedly, we found that the global transcriptional landscape was dominated by a pervasive Doxycycline-Responsive Gene Signature (DRGS), a systemic stress response that effectively masked transgene-specific effects at the aggregate gene level. We successfully decoupled the inducer artifact to isolate genuine trans-regulatory events. We identified six lncRNAs (DANCR, FENDRR, LINC00667, LINC00847, LNCPRESS1, and PNKY) that regulate specific transcript expression in trans, but we did not detect cis-regulatory effects from the engineered transgenes. Among these, LINC00847 showed a spatially biased trans-regulatory pattern, with its targets enriched on chromosome 19 and associated with RNA metabolism machinery. Overall, by disentangling pervasive Dox-induced stress from genuine regulatory signals, our work establishes a critical methodological baseline for the field while identifying temporally regulated trans targets for future mechanistic studies.

## Results

### Establishment of an Inducible lncRNA overexpression platform in hiPSCs

Based on our previous study on Firre, we learned that if a lncRNA can regulate gene expression in trans, the direct targets will be seen at early time points after lncRNA induction. To test the universality of this principle and identify the effects of specific lncRNAs on gene expression and epigenetic regulation in the context of pluripotency, we engineered a doxycycline-inducible expression platform in hiPSCs by integrating the Tet-On expression cassette and LoxP sites (Araki et al. 2002) into the H11 “safe-harbor” locus(Zhu et al. 2014) on chromosome 22 (Figure 1A).

**Figure 1.**
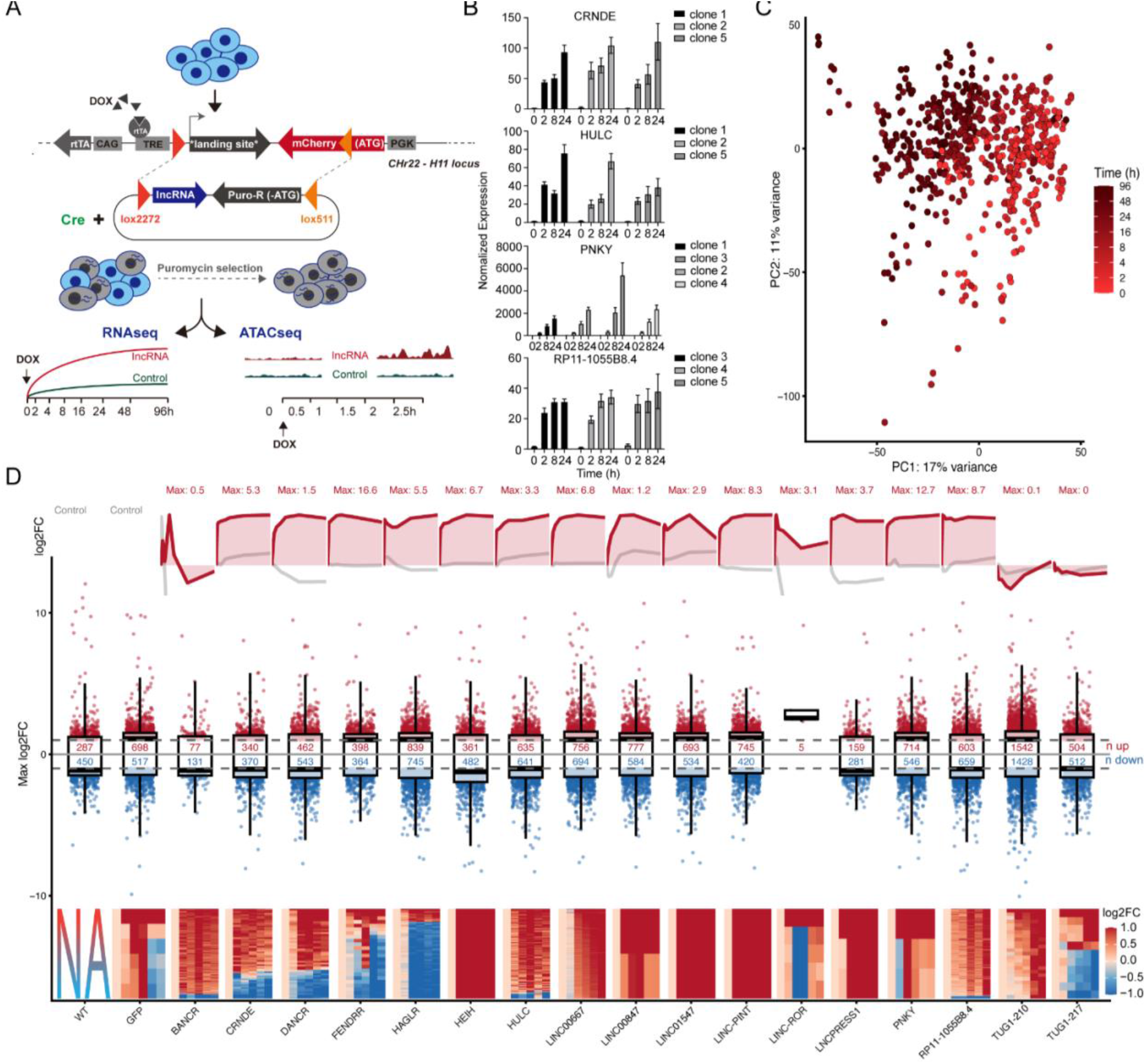
High-resolution RNA-seq and ATAC-seq atlas of induced lncRNA overexpression in human iPSCs. (A) Schematic diagram of the experimental design. The Tet-On inducible expression cassette containing each lncRNA gene was integrated into the H11 locus through Cre-LoxP recombination (Lox2272/Lox511), followed by puromycin selection. The bottom timeline shows the high-density sampling strategy for RNA-seq (0–96 hours) and ATAC-seq (0–2.5 hours) after doxycycline induction. (B) RT-qPCR verification of induced expression. The bar graph shows the normalized expression levels of four representative lncRNAs in different clones and time points. The data are presented as mean ± standard deviation (n=3 independent RNA isolations). (C) PCA of temporal transcriptomes. The PCA plot of all RNA-seq samples, colored by induction time (0–96 hours). (D) Global view of transcription and chromatin dynamics. Top: Temporal induction profiles detected by RNA-seq, showing the log2FC (log_2_ fold change) of specific lncRNAs relative to 0h within a 96-hour window. “Max” indicates the observed maximum log2FC. Middle: Distribution of significantly differentially expressed genes (DEGs) in all cell lines. The box plot shows the distribution of the maximum log2FC (relative to 0h) of all significant DEGs. The central line represents the median; the box boundaries represent the 25th and 75th percentiles; the whiskers extend to 1.5 times the interquartile range (IQR). Red and blue numbers represent the number of upregulated and downregulated genes that meet the significance criteria (padj< 0.01, baseMean>20, maximum |log2FC|>1 at any time point, and |log2FC|>0.58 at 24, 48, 96 hours). Bottom: Heatmap showing the signal intensity of significantly different ATAC-seq peaks in the early induction stage (0–2.5 hours). Columns represent time points (0, 0.5, 1, 1.5, 2, and 2.5 hours). NA indicates that ATAC-seq data for the WT cell line were not collected.

Using this platform, we generated 17 lncRNA-expressing cell lines (Table 1) and one control cell line expressing mRNA for Green Fluorescent Protein (GFP, Supplementary Figure S1C). These 17 lncRNA candidates were selected based on existing literature demonstrating robust in vitro or in vivo phenotypes upon genetic ablation or silencing. For loci with multiple splice variants, we cloned the specific isoform previously validated in the respective functional studies. In cases where isoform-specific information was unavailable, we selected the most abundantly expressed transcript isoform based on human tissue-wide RNA-seq data from the GTEx consortium V8 release (Consortium 2020). For each cell line, we picked at least 5 puromycin-selected single-cell colonies as biological replicates. To exclude chromosomal abnormalities that may be introduced by the genome editing process and subsequent resistance screening, we selected the GFP control cell line for G-band karyotype analysis. Results showed a normal karyotype (Supplementary Figure S1A).

**Table 1.**
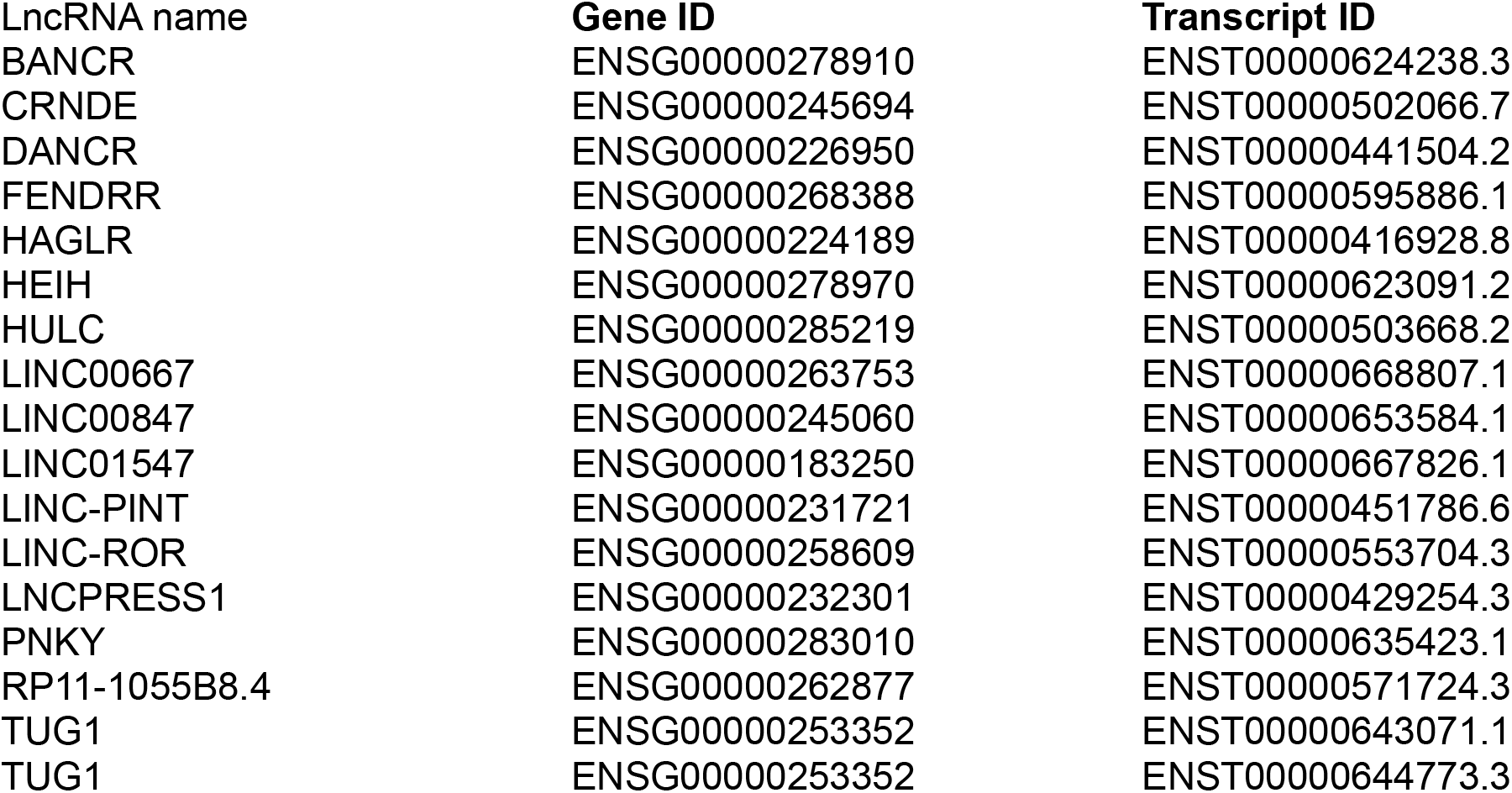
LncRNA Selected.

| lncRNA name | Gene ID | Transcript ID |
| --- | --- | --- |
| BANCR | ENSG00000278910 | ENST00000624238.3 |
| CRNDE | ENSG00000245694 | ENST00000502066.7 |
| DANCR | ENSG00000226950 | ENST00000441504.2 |
| FENDRR | ENSG00000268388 | ENST00000595886.1 |
| HAGLR | ENSG00000224189 | ENST00000416928.8 |
| HEIH | ENSG00000278970 | ENST00000623091.2 |
| HULC | ENSG00000285219 | ENST00000503668.2 |
| LINC00667 | ENSG00000263753 | ENST00000668807.1 |
| LINC00847 | ENSG00000245060 | ENST00000653584.1 |
| LINC01547 | ENSG00000183250 | ENST00000667826.1 |
| LINC-PINT | ENSG00000231721 | ENST00000451786.6 |
| LINC-ROR | ENSG00000258609 | ENST00000553704.3 |
| LNCPRESS1 | ENSG00000232301 | ENST00000429254.3 |
| PNKY | ENSG00000283010 | ENST00000635423.1 |
| RP11-1055B8.4 | ENSG00000262877 | ENST00000571724.3 |
| TUG1 | ENSG00000253352 | ENST00000643071.1 |
| TUG1 | ENSG00000253352 | ENST00000644773.3 |

Next, we tested the induction efficacy for an average of 2 clones at 0, 2, 8, and 24 hours by RT-qPCR. The results showed that Dox treatment induced significant upregulation of all lncRNA transgenes (Figure 1B, Supplementary Figure S1B). 14 out of 17 lncRNAs were largely induced at 2 hours after dox induction and continued to increase at 24 hours. Next, we combined high-temporal-resolution RNA-seq (0, 2, 4, 8, 16, 24, 48, and 96 hours, 3 replicates) and ATAC-seq (every 30 minutes from 0 to 2.5 hours, 3 replicates), which enabled us to capture the dynamics of both gene expression programs and chromatin accessibility landscapes after lncRNA induction.

In total, 663 RNA-seq datasets and 341 ATAC-seq datasets were generated, which provided a solid data foundation for analyzing regulatory events of lncRNAs at high temporal resolution.

### Global transcriptional and chromatin dynamics unfold along a temporal trajectory

To reveal the dynamic changes in gene expression and chromatin accessibility after lncRNA induction, we performed principal component analysis (PCA) on all RNA-seq data. The result showed a clear global trajectory that correlated with doxycycline induction time (Figure 1C). To identify temporal dynamic features in gene expression and chromatin accessibility, we used likelihood ratio tests (Love et al. 2014) to compare the changes of each cell line at different induction time points relative to the 0-hour baseline. Significant differences were observed in induction profiles of the lncRNA transgenes (Figure 1D, Top). Some lncRNAs (such as LNCPRESS1, PNKY) showed sustained and strong upregulation. BANCR showed a dynamic pattern of first rising and then falling. Despite successful verification by qPCR (Supplementary Figure S1B), two isoforms of TUG1 showed no significant upregulation in RNA-seq.

In most cell lines, we observed widespread genome-wide transcriptional perturbations upon Dox induction (Figure 1D, middle). Notably, the different cell lines exhibited a similar magnitude of transcriptional effects, typically manifested as hundreds of differentially expressed genes (DEGs), with similar numbers of up-regulated and down-regulated genes (see Figure 1D for “n up” and “n down” markers). More importantly, numerous DEGs were detected even in GFP and WT control cell lines without specific lncRNA expression. This phenomenon strongly implied that the broad transcriptional changes observed may not be driven by specific lncRNAs, but rather reflect a systemic response to the inducer itself (Ahler et al. 2013).

Similarly, our ATAC-seq analysis revealed a very early chromatin state change (Figure 1D, Bottom). Within the first 2.5 hours after Dox induction, we detected significant changes in chromatin accessibility in all cell lines tested, although transcriptional changes in some downstream genes may not be fully established at this time point. These rapidly occurring chromatin remodeling events, together with subsequent transcriptomic perturbations, constitute a continuous process. This decoupling between large gene expression changes independent of the genotype (each lncRNA transgene clone) and the induction efficiency of each lncRNA transgene suggested a non-lncRNA-specific driver in this system, which is Dox treatment.

### Defining a transcriptional signature driven by doxycycline in hiPSCs

To uncouple genuine transgene-specific regulatory events from the Dox-induced background, we first needed to define this systemic baseline. Through cross-sample overlap analysis (Supplementary Figure S2A-B), we defined a stringent signature set consisting of 209 genes, named the doxycycline response gene signature (DRGS), that were significantly differentially expressed in at least 21 biological replicates (≥75%, p=0.004, binomial test, Supplementary Table 2). Supplementary analysis confirmed that the Dox-induced signature was also pervasive at the transcript levels (Supplementary Figure S2C-D). To rule out any artifacts and or determine if there were local “*cis*” effects caused by transgenic integration, we examined the insertion region at the H11 locus on chromosome 22(Mele and Rinn 2016). Within a 4 Mb wide window centered on the insertion site (± 2 Mb), no DRGS genes were found. This strict negative result ruled out the possibility that the transgenic insertion caused changes in gene expression. Moreover, this also indicated that the induction of the lncRNA from the H11 locus did not cause cis-regulatory events by the lncRNA insert or the act of transcription of that insert.

Beyond their high recurrence across samples, we next explored whether the temporal dynamics of these DRGS genes were uniformly conserved across all transgenes. Quantitative comparison revealed a remarkable level of synchronization; specifically, a sub-module of 43 genes was consistently dysregulated in approximately 90% of all tested biological replicates, highlighting the pervasive and uniform nature of this systemic response. Heatmap visualization further confirmed that the DRGS genes exhibit a striking, highly synchronized transcriptional trajectory across all cell lines and time points, completely independent of the specific lncRNA transgene being induced (Figure 2A).

**Figure 2.**
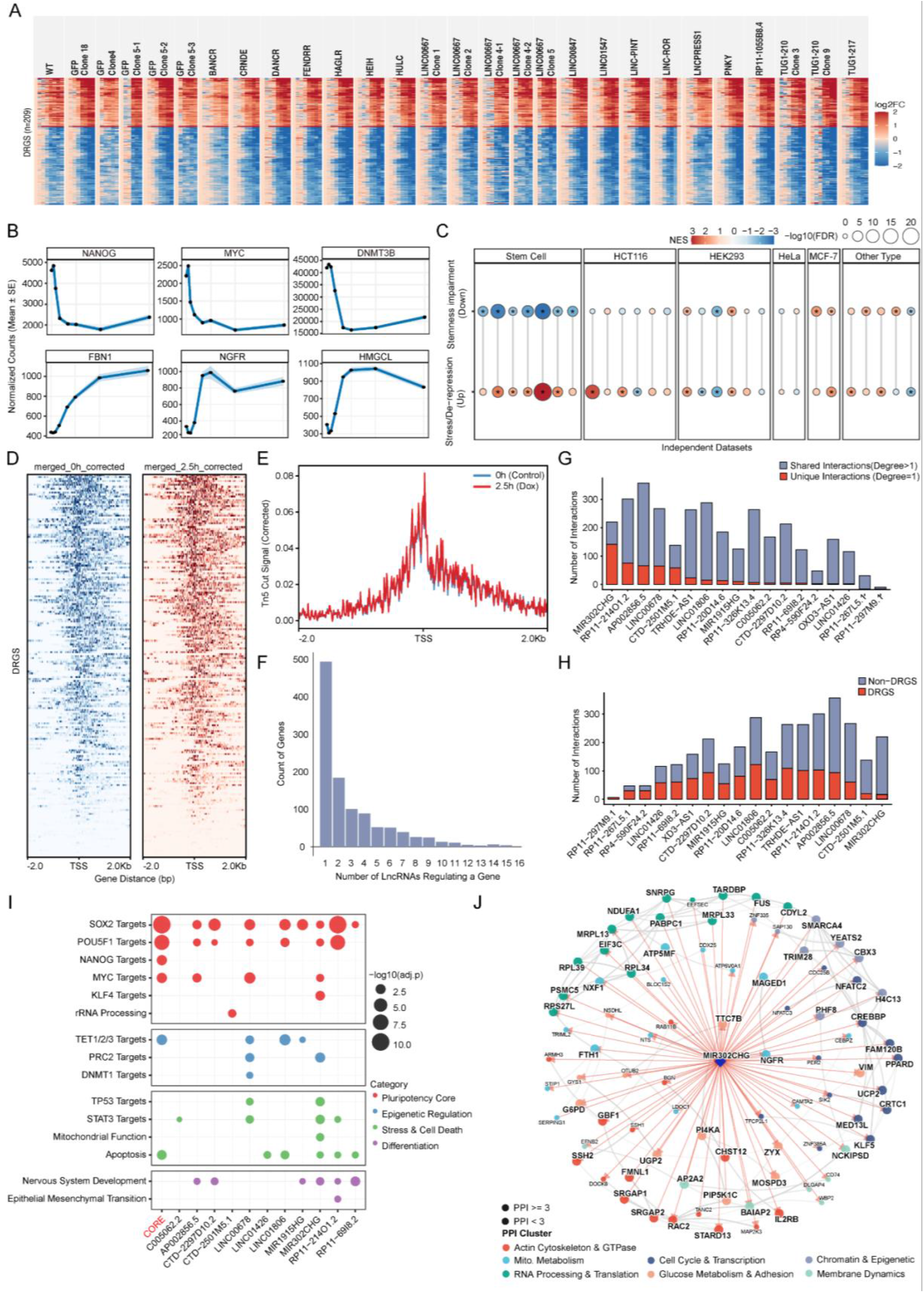
Identification of core transcriptional features of Dox response. (A) Time consistency of 209 core Dox response genes. The heatmap shows preexisting expression differences and subsequent expression changes (average log2FC relative to 0h) of the 209 core genes (defined as recurrence rate >= 75%) in all cell lines at time points 0, 2, 4, 8, 16, 24, 48, and 96 hours. (B) Ribbon plot of temporal expression profiles of representative genes in DRGS (vst-normalized counts, mean +/-standard error across 28 biological replicates). Pluripotency core factors (NANOG, MYC, DNMT3B) are significantly downregulated, while genes related to stress, metabolism, and differentiation (NGFR, FBN1, HMGCL) are significantly upregulated (padj < 0.01, baseMean > 20, maximum |log2FC| > 1 at any time point, and |log2FC| > 0.58 at 24, 48, 96 hours). (C) Meta-analysis of GSEA across 28 independent public datasets. DRGS is split and defined as “dysregulation of the stemness” and “stress/de-repression” modules. Bubble color represents the normalized enrichment score (NES), and size represents significance. The results show that the “stress (up)” module is broadly detectable across various cell types (red), and the “stemness (down)” module shows consistent downregulation only in stem cells (blue). Asterisks indicate false discovery rate FDR < 0.05. (D-E) Preexisting chromatin accessibility at promoter regions of the 209 core Dox response genes. (D) Heatmap of Tn5 cleavage signal at 0 hours and 2.5 hours, and (E) average signal contour map around the transcription start sites (TSSs) of the 209 core Dox response genes. (F) Scale-free network topology derived from integrated inference. (G) Specificity and redundancy. Stacked bar charts show the specificity (degree = 1, red) and shared (degree > 1, blue) node numbers of each DRGS lncRNA. (H) Network composition. Stacked bar charts quantify the proportion of DRGS genes (red) and non-DRGS genes (blue) in each lncRNA prediction network. (I) Bubble plots summarize the enrichment of pathways and transcription factor targets. “CORE” shared module represents the nodes connected to at least 5 DRGS lncRNAs. (J) Visualization of inferred subnetworks of MIR302CHG. The inferred associated genes are mapped to the STRING protein-protein interaction (PPI) database. Nodes are colored according to functional clusters (defined by STRING PPI), and red arrows represent directed associations inferred by DBN.

Given the robust and temporal properties of DRGS, we wanted to understand the biology underlying this exquisitely precise regulatory program. The downregulated protein-coding genes predominantly featured core components of the pluripotency network (e.g., NANOG, MYC) and key epigenetic regulators governing stemness (e.g., DNMT3B). Conversely, the upregulated genes were strongly linked to cellular stress, metabolism, and differentiation pathways, exemplified by the nerve growth factor receptor NGFR, the extracellular matrix protein FBN1, and the metabolic enzyme HMGCL (Figure 2B, Supplementary Figure S3A-B).

To determine whether the DRGS represents a universal phenomenon or an iPSC-specific artifact, we performed a meta-analysis of public transcriptomic data sourced from the Gene Expression Omnibus (Barrett et al. 2013). Specifically, we curated 28 independent datasets of Dox-treated human cell lines, requiring each study to have at least three biological replicates. We partitioned the 209-gene DRGS into up-regulated and down-regulated subsets and performed Gene Set Enrichment Analysis (GSEA) across all 28 datasets. We found that 21 out of the 28 datasets (75%) exhibited significant enrichment (FDR < 0.05) for at least one of the DRGS modules (Fig. 2C, Supplementary Figure S4). While the enrichment patterns displayed expected cell-type variations—most notably, all seven independent stem cell datasets exhibited significant and directionally consistent enrichment with our data—the signature was broadly detectable across diverse somatic and cancer cell types (e.g., HCT116, HeLa, HEK293). Collectively, this analysis confirmed that the DRGS is not an isolated experimental artifact, but rather a highly reproducible and widespread transcriptional response to doxycycline exposure across human cell lines.

### Epigenetic priming facilitates rapid transcriptional response

To understand the chromatin accessibility upstream from this rapid transcriptional upheaval, we analyzed high-temporal-resolution ATAC-seq data (0-2.5h, every 0.5h), where peaks represent chromatin accessibility. Despite the drastic remodeling of the transcriptome, we did not detect consistent differential peaks across cell lines, in contrast to the consistent RNA expression changes. Next, we analyzed the ATAC-seq signals 2kb upstream and downstream of the transcription start sites (TSS) of DRGS genes. Results indicated that these TSS regions were already highly accessible before Dox induction (0h), and after 2.5 hours of induction, the chromatin accessibility near the TSS remained virtually unchanged relative to the baseline, despite the massive onset of gene expression changes (Figure 2D-E). This demonstrates that DRGS genes reside in a “primed” state, allowing Dox to trigger immediate transcriptional bursting without the prerequisite of extensive de novo chromatin remodeling.

### Examination of lncRNA roles in DRGS

We noted that 18 endogenous lncRNAs are embedded within the DRGS signature. To understand their roles in the cellular response to Dox, leveraging the scale of our time-series data, we applied a Random Forest-guided ensemble Dynamic Bayesian Network (RF-DBN) strategy. Firstly, the RF regression model (Breiman 2001; Huynh-Thu et al. 2010) was used to screen out the Top 2000 feature genes that are most critical for predicting lncRNA dynamics. The model showed high prediction accuracy (average R^2^ = 0.85, Supplementary Figure S5A). Subsequently, to ensure the robustness of inference, we implemented a rigorous subspace resampling (Ho 1998; Marbach et al. 2012) strategy: The partitioning of these 2000 genes into 960 overlapping feature subspaces of 50 genes each ensured that each gene was independently evaluated in at least 20 different network contexts. Within each subspace, we further performed structure learning using DBN algorithms (Husmeier 2003; Scutari 2010) combined with 25 Bootstrap resamplings (Efron and Tibshirani 1994). Finally, to minimize false positives, we applied the most stringent filtering criteria: only those robust regulatory relationships that had a detection frequency of 1.0 in all Bootstrap iterations were retained.

The global network thus constructed shows a typical scale-free topology (BarabáSi and Albert 1999), with most genes in the network only associated with one or a few DRGS lncRNAs (Figure 2F). The global consistent scores, defined as the normalized bootstrap detection frequency across all evaluated feature subspaces (groups), exhibited a top-heavy distribution, which further confirmed the statistical robustness of these inferred relationships (Supplementary Figure S5B). These genes were selected from the DRGS-associated regulatory network inferred from the 209 recurrent Dox-responsive genes and the endogenous DRGS lncRNAs. By quantifying the degree of interaction sharing (Figure 2G) and network composition (Figure 2H), we observed a division-of-labor pattern among the lncRNAs. The results are represented by MIR302CHG, which has the most unique interactions and the fewest interactions with DRGS genes, implying it may play a role in functional spreading during Dox response.

To further understand the role of these lncRNAs in stress response, we did enrichment analysis on “MsigDB Hallmark 2020(Liberzon et al. 2015)”, “Reactome 2022(Gillespie et al. 2022)”, and “TF Perturbations Followed by Expression(Kuleshov et al. 2016)” terms (Figure 2I). The results demonstrated that the “CORE” shared module (genes associated with at least five lncRNAs) is highly enriched for targets of key pluripotency factors (SOX2, POU5F1, NANOG, MYC), and DNA demethylases (TET1/2/3), alongside apoptosis-related genes. Unlike the shared module, the target genes of specific epigenetic silencers (e.g., PRC2, DNMT1) and distinct developmental fates were exclusively enriched in the subnetworks of specific “specialist” lncRNAs. For instance, RP11-69I8.2 is specifically linked to nervous system development, whereas MIR302CHG is linked to mitochondrial function. To determine whether the statistically inferred dependencies reflect known biological functional modules, we mapped the DBN-derived subnetwork nodes of MIR302CHG (Figure 2J) onto the protein-protein interaction (PPI) network constructed based on the STRING database (Szklarczyk et al. 2023). Visualization revealed that the DBN-inferred genes did not appear as isolated entities, but rather clustered into established interaction networks.

### Transcript-level analysis identifies lncRNA-specific targets beyond the Dox response

Having established that the DRGS dominates gene-level variance across all cell lines, we next sought to identify authentic regulatory effects of individual lncRNA transgenes. Since gene-level quantification aggregates all transcripts from a given locus, it masks regulatory events such as changes in promoter selection, exon usage and differential polyadenylation, mechanisms recognized as primary targets of lncRNA regulation (Kopp and Mendell 2018; Statello et al. 2021; Mattick et al. 2023). We therefore shifted our framework to the transcript/isoform level, which provides resolution to distinguish transcript-specific regulation signals.

To distinguish transgene-specific effects, we developed a stringent filtering strategy combining multiple orthogonal criteria. First, we performed pairwise correlation analyses (Pearson and Spearman) between the expression of all the transgenes and all potential target transcripts with sufficient abundance (>100 counts) across time points. This approach allowed us to capture both linear and monotonic non-linear associations. We then applied a five-step filtration process to identify robust regulatory relationships: (1) high correlation (Pearson or Spearman > 0.75, padj < 0.01; Supplementary Figure S6A); (2) specific induction, where the target gene showed significant higher expression (> 2 log fold change) in the lncRNA-induced condition compared to both GFP and WT controls; (3) a significant temporal trajectory (absolute slope > 0.15, P < 0.05); and (4) a distinct expression curve deviation relative to GFP (> 5 log difference) and WT (> 5 log fold change) controls, using the statistical curve-deviation threshold shown in Figure 3A. (5) Finally, candidate targets underwent manual visual inspection to verify trajectory consistency and rule out fitting artifacts. Crucially, for lncRNAs represented by multiple clones (e.g., LINC00667 and TUG1-210), clones were analyzed independently, and only targets passing criteria 1-4 across all biological replicates were retained.

**Figure 3.**
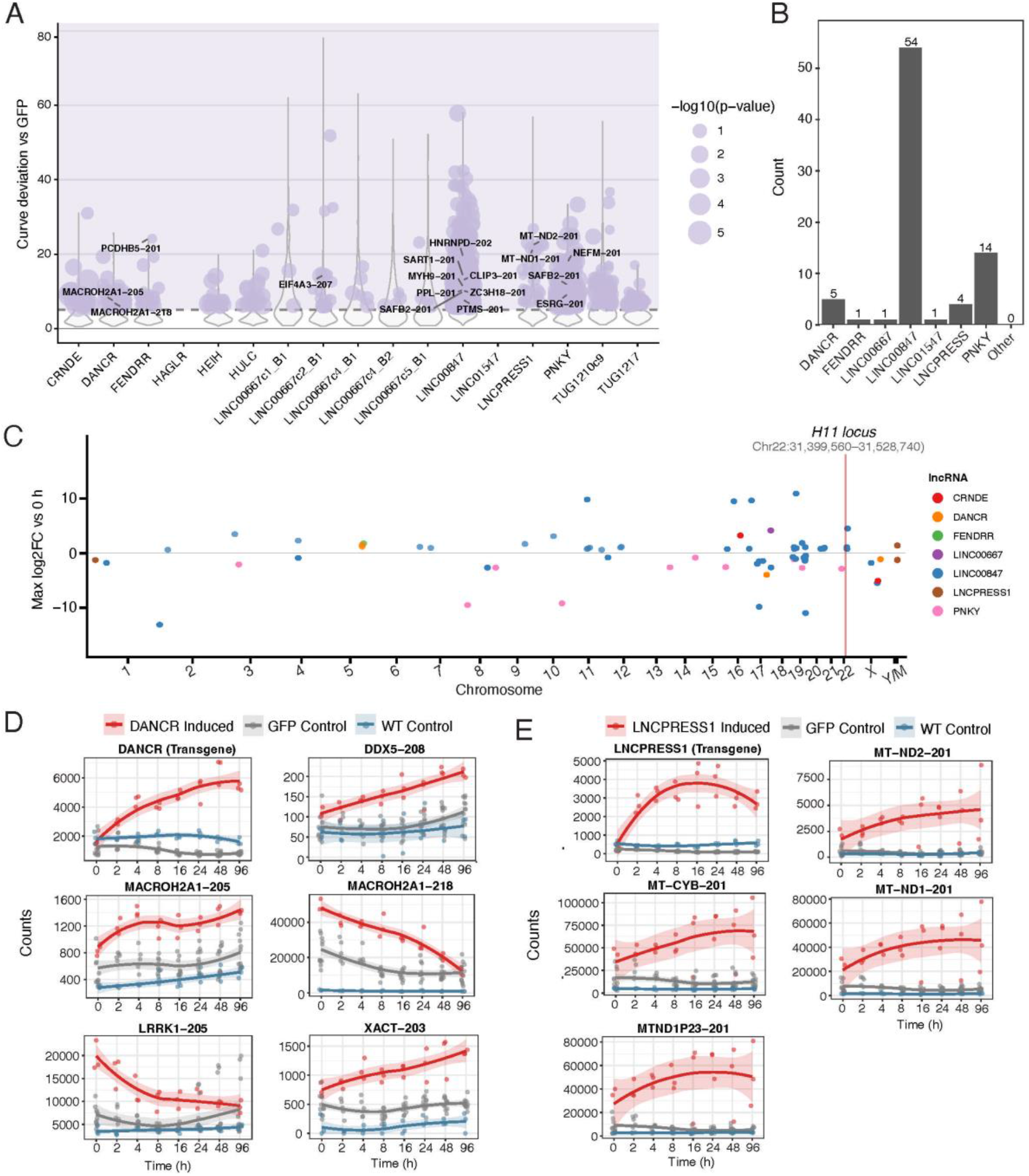
Identification of transcript-level lncRNA-specific trans-targets. (A) Curve deviation versus GFP control across transgene libraries. Violin plots display the distribution of curve deviation scores, quantifying expression trajectory differences between transgene-induced and GFP control conditions for each lncRNA library. Purple circles mark transcript pairs that pass the curve-deviation threshold with significant p-values; circle size represents -log10(p-value). The shaded region indicates the filtering threshold for transgene-specific effects. Selected high-confidence targets are labeled (MACROH2A1 isoforms, PCDHB5-201, EIF4A3-207, MT-ND1/ND2-201, and others). This metric identifies targets whose temporal profiles are specifically altered by transgene induction beyond the doxycycline response. BANCR, HAGLR, and LINC01547 did not yield targets that passed the correlation threshold of 0.75 and padj < 0.01 and therefore are not represented in the target-count summary. (B) Distribution of transgene-specific transcript pairs across lncRNA libraries. The bar chart shows the number of high-confidence target transcripts identified for each lncRNA transgene after applying stringent filtering criteria. LINC00847 showed the highest number of putative regulatory targets (n = 54). (C) Scatter plot visualizing the genomic loci and expression magnitude (Max log2FC relative to 0 h; y-axis) of all 78 robustly identified lncRNA targets across the human genome (x-axis, concatenated chromosomes 1 through Y and mitochondrial DNA). Dots are color-coded by the specific inducing lncRNA transgene. A solid vertical red line demarcates the precise H11 integration locus on chromosome 22; all dots shown near the line are over 2 Mb away from the insertion site on chr22. Notably, zero high-confidence targets were detected in the immediate genomic vicinity of the H11 locus, confirming the absence of local cis-regulatory artifacts. Instead, the specific targets exhibit a widespread trans-chromosomal distribution, with a prominent clustering of LINC00847 targets (blue dots) specifically enriched on chromosome 19. (D-E) Temporal expression dynamics and target transcripts of DANCR (D) and LNCPRESS1 (E). Time-course plots show raw counts (y-axis) across eight timepoints (x-axis) for the transgene (top left) and high-confidence targets. Each point represents an individual biological replicate. Red: transgene-induced (DANCR or LNCPRESS1) library; gray: GFP control; blue: WT control. Lines represent LOESS-smoothed trends.

This stringent strategy identified 78 high-confidence lncRNA-target pairs across 6 transgenes (Figure 3B). Importantly, none of these 78 transcripts overlapped with the generic doxycycline signature, and unlike the DRGS genes, which showed uniform patterns, these transgene-specific targets displayed individualized expression trajectories. Notably, several induced lncRNAs, including BANCR, CRNDE, FENDRR, HAGLR, HEIH, HULC, LINC-PINT, LINC-ROR, and RP11-1055B8.4, did not yield putative targets passing these strict criteria. To confirm that our system evaluates pure trans-acting capacities, we visualized the genome-wide distribution of these lncRNA targets. Spatial genomic analysis revealed an absence of associated transcripts within a conservative window surrounding the H11 integration site on Chromosome 22 (Figure 3C, Supplementary Figure S7).

Each lncRNA showed a distinct regulatory signature (Figure 3D-E, Supplementary Figure S6B-D). DANCR, a lncRNA previously implicated in differentiation antagonism and recently associated with development in human and zebrafish(Jones et al. 2025), exhibited 5 high-confidence targets, including two isoforms of MACROH2A1 (MACROH2A1-205 and MACROH2A1-218), which encode a histone variant involved in transcriptional repression and X-chromosome inactivation (Figure 4D). The coordinated upregulation of these histone variant transcripts suggests DANCR may influence chromatin state in human pluripotent stem cells. Additional DANCR targets included DDX5-208 (a DEAD-box RNA helicase), XACT-203 (X-active coating transcript), and LRRK1-205 (leucine-rich repeat kinase 1). Notably, LNCPRESS1 demonstrated specific upregulation of multiple mitochondrial-encoded transcripts, including MT-ND1-201, MT-ND2-201, MTND1P23-201, and MT-CYB-201 (Figure 4E). These targets encode subunits of the mitochondrial respiratory chain complexes (NADH dehydrogenase Complex I subunits ND1 and ND2, and cytochrome b of Complex III). LNCPRESS1 was first described as a p53-regulated transcript(Jain et al. 2016), highly expressed in human ESCs and repressed by p53 during differentiation, coupled with metabolic rewiring. PNKY, a lncRNA previously characterized as a regulator of cortical neurogenesis(Ramos et al. 2015; Andersen et al. 2019), showed 14 putative targets enriched for neuronal differentiation markers and developmental regulators (Supplementary Figure S6B). Targets included NEFM-201 (neurofilament medium polypeptide, a marker of mature neurons), ESRG-201 (embryonic stem cell-related gene), LRRK1-205, and SAFB2-201 (scaffold attachment factor B2, involved in RNA processing and transcriptional regulation).

**Figure 4.**
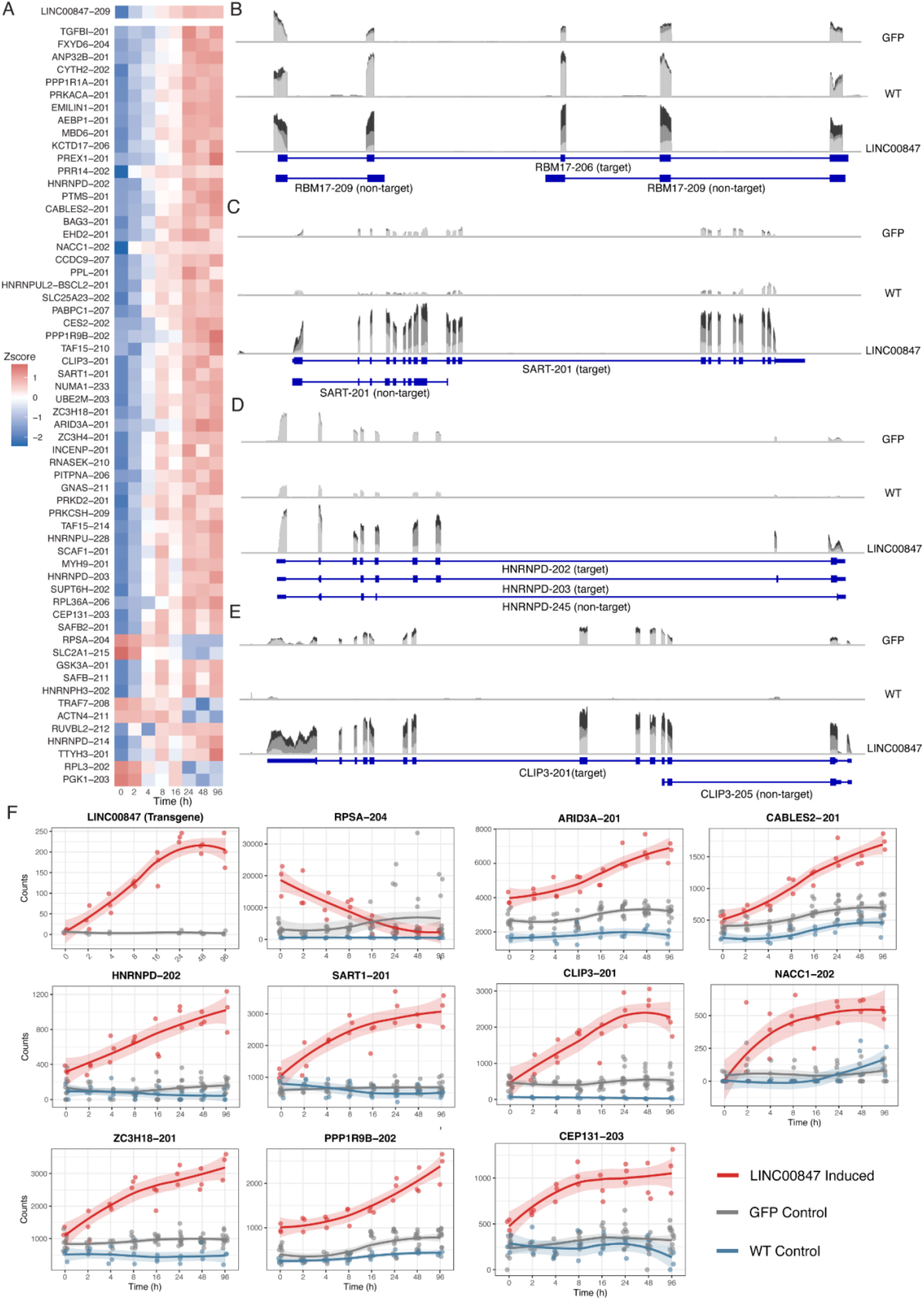
High-resolution transcriptomic profiling reveals trans-regulation by LINC00847. (A) Heatmap showing z-score normalized expression patterns of all 54 LINC00847-specific targets across the timepoints. Each row represents a transcript. Columns represent timepoints. Color scale represents z-score: blue (low expression), white (baseline), red (high expression). LINC00847 transgene (LINC00847-209) is shown at the top, followed by all identified targets. (B-E) Genome browser tracks visualizing isoform-specific trans-regulation by LINC00847. RNA-seq read coverage tracks across the time course (0h in light gray, 8h in dark gray, 24h in black) are displayed for representative high-confidence target transcripts: (B) RBM17-206, (C) SART1-201, (D) HNRNPD-202, and (E) CLIP3-201. For each panel, the top two tracks represent the control conditions (GFP and WT), while the bottom track represents the LINC00847-induced condition. The blue schematic at the bottom of each panel denotes the exon-intron structure of the specifically regulated transcript isoform and non-target transcript. Upon LINC00847 overexpression, these exonic regions exhibited increased read abundance, whereas the corresponding loci in both GFP and WT controls remained largely unchanged. (F) Temporal expression dynamics of LINC00847 and representative target transcripts. Time-course plots show raw counts (y-axis) across eight timepoints (x-axis) for LINC00847 transgene (top left) and 12 selected high-confidence targets. Each point represents an individual biological replicate. Red: LINC00847-induced gene; gray: GFP control; blue: WT control. Lines represent LOESS-smoothed trends. Target transcripts displayed altered expression in the LINC00847-induced condition compared to both control conditions, with minimal or no induction observed in GFP or WT controls.

To better understand these trans-regulatory events, we characterized the biotypes of 78 high-confidence target transcripts. The majority were mRNAs (n=53), whereas 10 were annotated as lncRNAs. The remaining 14 targets contained non-cononical or alternative RNA processing variants, including transcripts that underwent nonsense-mediated decay (NMD, n=4) and intron retention (n=6). We next investigated whether trans-regulation of these targets occurs as an isolated event or as part of a broader regional transcriptional activation. We examined a ±1 Mb window around each of the 78 target loci to see if any neighboring genes were co-activated or co-repressed. We found that the apparent spatial clustering was almost entirely driven by variable isoforms of the same gene (e.g., HNRNPD, MACROH2A1) or transcripts within the compact mitochondrial genome. In addition, only four pairs of target genes were spatially adjacent, and all mapped to the gene-dense chromosome 19. Among the 10 lncRNAs, nine were significantly downregulated, and we did not observe any co-repression or co-activation of other genes within their ±1 Mb vicinity, which might occur if these lncRNAs functioned in cis, as shown previously for other cis-acting lncRNAs(Unfried and Ulitsky 2026).

### LINC00847 regulates targets in a specific chromosome in trans

Among the active lncRNAs, LINC00847 emerged as having 54 putative targets, the highest number of the tested lncRNAs (Figure 3D). Heatmap visualization confirmed that these specific transcripts displayed highly individualized expression trajectories coupled with LINC00847 induction, distinct from the uniform pattern of the DRGS (Figure 4A). To visually validate these isoform-specific variations, we examined the sequencing read coverage using BigWig tracks (Figure 4B-E). The alignment profiles showed that LINC00847 induction selectively alters the read abundance of specific target isoforms (e.g., RBM17-206 and CLIP3-201), while non-target variants transcribed from the same loci remain unaffected. For representative targets, the tracks displayed pronounced, time-dependent alterations in read abundance at specific exonic regions following LINC00847 induction (Figure 4F). Crucially, these genomic regions remained unchanged in both wild-type and GFP control cell lines across all time points. This direct physical evidence confirmed that the observed transcript-level modulations were strictly LINC00847-specific.

We next annotated the biological functions of the 54 unique target transcripts (of 51 genes). This revealed a pronounced bias toward RNA metabolism. Specifically, 17 out of 51 target genes encode proteins centrally involved in the RNA metabolism machinery. Notable examples include major heterogeneous nuclear ribonucleoproteins (HNRNPU, HNRNPD, HNRNPH3), core spliceosome-associated factors (SART1, RBM17), and the polyA-binding protein PABPC1. Utilizing the ∼16,500 actively expressed genes in our hiPSCs as the background, this represents a 5-fold enrichment over random expectation (p=1.52e-08, hypergeometric test), indicating a significant functional association with the cellular RNA processing machinery.

Alongside this functional coherence, mapping the genomic distribution of these targets revealed a clear spatial bias. Although LINC00847 is transcribed from Chromosome 22, of its 54 trans-target transcripts, 16 are localized on Chromosome 19, but not clustered in proximity (Figure 3E). Considering that only 5.9% of the expressed background transcripts in our dataset are located on Chromosome 19, observing 29.6% of the targets on this single chromosome represents a significant 4.24-fold enrichment (p =1.19e-06, hypergeometric test).

## Discussion

Many lncRNAs have been found to function in cis, including well-studied examples such as XIST(Penny et al. 1996) and enhancer-like lncRNAs such as HOTTIP(Wang et al. 2011) and PVT1(Cho et al. 2018), where transcriptional activity or RNA products regulate local chromatin and gene expression. By contrast, trans activity has been more elusive because it requires separating primary RNA-mediated effects from secondary responses and from the native genomic context of the lncRNA locus. Here, we tested this directly by evaluating the regulatory capacity of 17 lncRNAs in hiPSCs using a high-resolution, time-resolved ectopic expression system. This design isolates lncRNAs from their native genomic background, similar to evaluating the trans-acting effects of exogenously delivered RNA therapies (Pang et al. 2026) and identified six lncRNAs with specific trans-regulatory effects.

The most prominent example in our study is LINC00847, which yielded the highest number of high-confidence trans-targets (n = 54 transcripts, mapping to 51 unique genes). Although transcribed from chromosome 22, 16 out of the 54 specific trans-targets of LINC00847 are significantly enriched on chromosome 19 (P = 1.413e-07). Furthermore, 17 out of 51 of its unique gene targets encode proteins functionally concentrated within the RNA processing machinery (e.g., SART1, HNRNP families, and ZC3H families).

This spatial and functional bias warrants mechanistic consideration. The targeting of an entire chromosome in trans evokes the classic model of XIST, which utilizes 3D genome conformation to spread across and silence the X chromosome in cis (Engreitz et al. 2013). Emerging evidence suggests that even this master cis regulator can localize in trans to autosomes under specific contexts (Dror et al. 2024; Weigel 2024). Another example is lncRNA Firre. It has been shown to mediate inter-chromosome interaction, anchoring multiple trans-chromosomal loci to specific nuclear compartments (Hacisuleyman et al. 2014). These observations suggest that some lncRNAs may participate in cross-chromosomal communication from one chromosome to another. Chromosome 19 is unique in the human genome: it is the most gene-dense chromosome; it accounts for only 2% of the human genome but harbors around 1500 genes, with an exceptional density of Alu repeats and clusters of genes encoding RNA-binding proteins and splicing factors (Chen et al. 2002; Grimwood et al. 2004; Grover et al. 2004; Lukic et al. 2014; Gosztyla et al. 2024). Although the mechanism and the directness of LINC00847 preferentially associate with chromosome 19 remain to be tested.

Meanwhile, the spatial and functional specificity of LINC00847 that we observed extends the existing biological paradigm of LINC00847. To date, the existing literature has characterized LINC00847 mainly in the context of oncology, identifying it as an oncogenic driver in malignant tumors such as non-small cell lung cancer, pancreatic cancer, and melanoma (Li et al. 2021a; Hao et al. 2024; Jiang et al. 2024; Liu et al. 2024). Mechanistically, these studies have primarily modeled LINC00847 as a cytoplasmic competing endogenous RNA (ceRNA) that acts as a molecular sponge to adsorb microRNAs. Our results add a nuclear, chromosome-biased trans-regulatory activity to these reported roles, while leaving open the possibility that LINC00847 has additional functions in other cellular contexts.

The majority of the lncRNAs that were robustly induced from the H11 safe harbor locus failed to yield detectable trans targets. There are several possibilities for this lack of observable trans-activity. It may primarily reflect the exquisite cell-type specificity inherent to lncRNA function.

LncRNAs frequently operate as modular scaffolds or guides, requiring precise stoichiometric interactions with specific RNA-binding proteins (RBPs) to execute their regulatory roles (Engreitz et al. 2016; Gil and Ulitsky 2020; Mattick et al. 2023). It is plausible that the pluripotency state lacks the requisite endogenous interacting partners for lncRNA action. For example, HULC and HAGLR are developmentally regulated and typically function in specific differentiated lineages. Consequently, when ectopically induced in hiPSC, these transcripts may be “orphaned” from their essential molecular machinery. As lncRNAs are notoriously tissue-specific and we only tested iPSCs, perhaps these lncRNAs would be active if they were expressed in a different chromosomal context or in a cell type other than iPSCs.

Beyond cell-type specificity, the absence of detectable trans-targets for some lncRNAs might relate to spatial constraints. Recent genetic models emphasize that numerous lncRNAs operate exclusively in cis to organize local topological association domains (TADs) or regulate proximal gene transcription through enhancer-like mechanisms (Engreitz et al. 2016; Gil and Ulitsky 2020). Furthermore, specific lncRNAs require native genomic contexts to form local nuclear condensates, such as the nucleolus (Savic et al. 2014). By ectopically expressing these lncRNAs at the H11 locus, our platform deliberately separates the lncRNA sequence from its native cis-regulatory environment. Finally, some lncRNAs may possess functions other than gene expression regulation (Clemson et al. 2009; Lee et al. 2016; Xing et al. 2017; Li et al. 2021b), which would inherently not be captured by transcriptomic profiling.

In summary, our study establishes DRGS as an important resource for decoupling general metabolic stress from specific regulatory signals in doxycycline-inducible systems. By leveraging high-resolution transcriptomic and ectopic expression platforms, we found that a minority of tested lncRNAs are sufficient to regulate transcript expression in trans, and local cis effects from the engineered transgenes were not detected. The identification of isoform-specific, chromosome-biased trans-regulation represented by LINC00847, including enrichment of targets on chromosome 19, provides a focused model for future mechanistic studies that connect 3D nuclear architecture with non-coding RNA biology.

### Limitations of this study

Although our orthogonal filtering strategy successfully separated trans-regulatory events from pervasive systemic effects of doxycycline, the current approach has certain limitations. Firstly, clonal variations may be introduced into stable cell lines selected by monoclonal screening. Although we isolated multiple independent clones for each lncRNA transgene and applied strict filtering criteria to identify reproducible targets, the cost of RNA-seq at intensive time points limited the testing of multiple biological replicates for most lncRNAs. Thus, the extent of clonal heterogeneity in the target genes of a given lncRNA remains to be evaluated. Second, our experiments were performed exclusively in hiPSCs. Given the strong cell and tissue specificity of lncRNA function, how these specific trans-regulatory targets change in other differentiated cell lineages remains to be determined. Finally, we captured transcriptome dynamics, but regulation of mRNA levels does not always result in alterations in protein abundance. Future studies employing functional proteomics will be essential to test whether the isoform-specific changes observed here are effectively translated into functional proteomic and phenotypic outcomes.

### Methods Cell Culture

Human-induced pluripotent stem cells (Applied StemCell, ASE-9211) were maintained in mTeSR Plus medium (STEMCELL Technologies, 100-0276) on the T25 flask coated with Geltrex (Gibco, A1413302). The cells were grown in a 5% CO2 incubator at 37 °C. The medium was changed daily. Upon reaching 70-80% confluency, the cells were passaged using Accutase (STEMCELL Technologies, 07922). For the first 24 hours after passaging, the medium was supplemented with 10 μM of Y-27632 (Watanabe et al. 2007) (STEMCELL Technologies, 72304).

### Plasmid construction

The gene fragments for the 17 lncRNAs we selected were synthesized by Twist Bioscience and cloned into the PCR-linearized pCR-XL-lox511-PURO-lox2271 vector using In-Fusion (Takara Bio) assembly according to the manufacturer’s instructions. Verification was performed by colony PCR and subsequent sequencing (Plasmidsaurus). Sequences of gene fragments and primers are listed in the supplement table.

### Stable cell lines generation

The Dox-inducible expression cassette and LoxP site were integrated into the H11 safe harbor locus on Chromosome 22 using the method described before (Chen-Tsai 2019). For transgene integration, the hiPSCs were dissociated using Accutase and seeded onto Geltrex-coated 6-well plates at a density of 3.5 × 10^^5^ cells per well. To enhance cell survival, the culture medium was supplemented with CEPT Cocktail (Chen et al. 2021) (MedChemExpress, HY-K1043). After 2 hours post-seeding, cells were transfected with 300 ng of Cre plasmid and 1800 ng of target gene plasmid using 7.5 μL of Lipofectamine Stem Transfection Reagent (Invitrogen, STEM00008). After 48 hours of transfection, the medium was replaced with fresh mTeSR Plus medium. When the transfected cultures reached 70-80% confluency, cells were passaged into Geltrex-coated 12-well plates in the presence of 10 μM Y-27632. On the following day, starting antibiotic selection by supplementing the medium with 200 ng/mL Puromycin (InvivoGen, ant-pr-1). The selection medium was refreshed daily until distinct single colonies appeared. Individual colonies were identified and manually picked under a microscope. Before single colony picking, cells were treated with 10 μM Y-27632 for 2 hours and washed twice with warm DPBS. Colonies were aspirated using a P200 pipette tip in 100 μL of PBS and transferred to Geltrex-coated 6-well plates containing mTeSR medium supplemented with 10 μM Y-27632. After 2 days of recovery, the selection pressure was increased to 500 ng/mL Puromycin. The medium was changed daily until the stable cell lines reached approximately 70-80% confluency, after which they were expanded and cryopreserved. For karyotype analysis, live cells were sent to ThermoFisher Scientific for G-banded karyotyping.

### Doxycycline Induction and qRT-PCR

Cells were treated with 1 μg/mL doxycycline (Sigma, D5207). Total RNA was extracted using TRIzol reagent (Invitrogen, 15596018) and Directzol RNA Miniprep kit (ZymoResearch, R2062). 500 ng of RNA was reverse transcribed into cDNA using SuperScript IV First-Strand Synthesis System (Invitrogen, 18091200). Quantitative real-time PCR was performed using LightCycler 480 SYBR Green I Master (Roche, 04707516001) on a CFX Connect Real-Time PCR Detection System (Bio-Rad). Relative gene expression levels were calculated using the 2^−ΔΔCt^ method (Livak and Schmittgen 2001), with GAPDH as the internal control. Primer sequences are available in the Supplementary Table 1.

### RNA-seq and ATAC-seq

For RNA-seq, libraries were prepared from ribosomal RNA-depleted total RNA and sequenced on an Illumina NovaSeq X plus platform using a paired-end 150 bp (PE150) configuration. The average sequencing depth was 100 million read pairs per sample. For ATAC-seq, libraries were sequenced on an Illumina NovaSeq X plus platform (PE150), with an average depth of 40 million read pairs per sample.

### RNA-seq Analysis

Quality control, read mapping, and quantification were performed using nf-core/rnaseq v3.14.0 (Ewels et al. 2020). The cleaned reads were aligned to the GRCh38.p13 human reference genome and quantified using GENCODE v38 annotations. The alignment process was performed using STAR v5.1.0 (Dobin et al. 2013), and the quantification process utilized Salmon v1.10.1 (Patro et al. 2017). For differential expression analysis, the raw count matrix was imported into R v4.3.2, then normalized and analyzed for differential expression using the DESeq2 v1.42.1 package (Love et al. 2014). For time-series data, the likelihood ratio test (LRT, design=∼time) was used to identify genes that significantly changed over time (cutoff: padj < 0.01, baseMean>20, maximum log2FoldChange >1, and log2FoldChange>0.58 at 24, 48, and 96 hours). To perform principal component analysis (PCA) and visualization, the variance stabilizing transformation (VST) was applied to the count matrix.

For transcript-level analyses, reads were pseudo-aligned and quantified with Salmon (Patro et al. 2017) against GENCODE v38 transcriptome reference using Gibbs Sampling Inferential replicates. Transcripts that were too similar to be distinguished were measured using Terminus (Sarkar et al. 2020) and removed from the analysis. Quant files were imported into R for downstream differential transcript expression and visualization analyses. Transcript-level differential expression was performed in edgeR v4 (Baldoni et al. 2024), modeling the time effects. Transcript-level time structure was summarized by WGCNA, and module–time associations were assessed using module eigengene correlations, with module enrichment and overlap with the core Dox response signature reported for modules most strongly associated with time. All processing and plotting were performed using tidyomics (Hutchison et al. 2024) and Isoformic (Mamede et al. 2025).

To evaluate whether transgene expression was associated with consistent transcript programs beyond the shared Dox response, we curated a “source-of-truth” map of each transgene to its intended ENST transcript(s) and, for each lncRNA counts file, computed pairwise correlations across samples between the transgene ENST expression vector and all other transcripts using a log2(count+1) transform. Both Pearson r and Spearman ρ were calculated, with two-sided P-values derived from an approximate t-statistic and Benjamini–Hochberg (BH) adjustment applied within each set; significant pairs were retained if either Pearson or Spearman nominal P-value < 0.01. To control for background correlations driven by Dox treatment and/or generic cassette effects, the same correlation procedure was run on GFP control count files for all target ENSTs, and transgene-associated pairs were filtered to remove any transcript pairs also significant in GFP. For candidates prioritized from the signed correlation set, we further quantified time-course specificity by extracting timepoints from sample identifiers, summarizing mean expression per timepoint for each condition (transgene line, GFP, WT), transforming to log2(mean+1), and computing curve deviation metrics (sum of absolute differences across shared timepoints and maximum absolute difference), together with global median/mean log-expression across time and linear trend slopes (lm(log expression ∼ time)) for interpretability. Resulting 1,304 pairs between induced lncRNA and correlated mRNA were then individually confirmed one by one.

### ATAC-seq Analysis

Quality control, read mapping, and quantification were performed using nf-core/atacseq v2.1.2. The cleaned reads were aligned to the GRCh38.p13 human reference genome using BWA-MEM v0.7.17-r1188 (Li and Durbin 2009), with the GENCODE v38 genome annotation file. Reads located in the ENCODE hg38 blacklist v2 region were removed. On the filtered and deduplicated alignment files, the broad peak calling was performed using MACS2 v2.2.7.1 (Zhang et al. 2008). The peaks from different biological replicates were merged using the built-in consensus strategy in the pipeline to generate a consensus peak set. Peak counts were calculated using featureCounts (Liao et al. 2014). Differentially accessible peaks were analyzed using DESeq2 v1.42.1 with LRT design= ∼time (cutoff: padj<0.05, baseMean>200, maximum log2FoldChange>1). Subsequently, chromatin openness profiles and heatmaps near the transcription start sites (TSS) were plotted using computeMatrix, plotProfile, and plotHeatmap in deepTools v3.5.1 (Ramírez et al. 2016). For TOBIAS footprint analysis (Bentsen et al. 2020), the TOBIAS ATACorrect module was first used to calculate and correct the sequence bias of the Tn5 transposase, generating a corrected signal file. The footprint score was calculated using the ScoreBigwig module. The BINDetect module was used to compare the TF binding differences before and after Dox treatment (2.5h vs 0h), generating a differential binding volcano plot and an aggregated footprint plot.

### Enrichment Analysis

The gene function enrichment analysis (GO Biological Process, KEGG) was conducted using the clusterProfiler v4.10.1 (Wu et al. 2021). The gene set enrichment analysis was employed to evaluate the enrichment of DRGS in public datasets. For the enrichment of transcription factor target genes and epigenetic modifications, Enrichr v3.4 (Kuleshov et al. 2016) and related databases (ENCODE, Epigenomics Roadmap) were utilized. All enrichment analyses were corrected for multiple hypothesis testing using the Benjamini-Hochberg method (FDR < 0.05).

### RF-guided ensemble DBN

In order to select the most predictive regulatory factors for the dynamic changes of lncRNAs from high-dimensional transcriptome data and eliminate collinearity noise, we adopted the random forest (RF) regression strategy. The analysis was performed using the caret v7.0-1 R package (Kuhn 2008) as a wrapper for the randomForest v4.7-1.2 package (Liaw and Wiener 2002). For each target lncRNA, a regression model was constructed. The response variable y was the normalized expression level (VST normalized counts) of the target lncRNA, and the predictor variable X included the expression levels of the remaining top 5000 highly variable genes, and the induction time points, to capture the temporal dependence. To prevent the model from overfitting the batch effects of specific cell lines and ensure generalization ability, we implemented strict 5-fold grouped cross-validation. Using the groupKFold function, we defined biological replicates as grouping variables. The model parameters were set as ntree=500 (number of decision trees), mtry (number of variables randomly sampled at each split) was set to be searched within the range of 50 to 500 (step size of 50) to find the optimal balance point between bias and variance. The importance of the feature genes was sorted based on %IncMSE (increase in mean squared error), and the top 2000 genes were selected for subsequent network inference.

To accurately infer the directed dependency relationships between genes and overcome the computational challenges of high-dimensional data, we developed an integrated dynamic Bayesian network (Ensemble DBN) strategy, implemented based on the bnlearn v4.9 R package (Scutari 2010). Drawing on the random subspace method (Ho 1998), we divided the 2000 selected feature genes into 960 overlapping feature subspaces. Each subspace contains 50 genes, generated through random sampling, ensuring that each gene is independently evaluated in at least 20 different network contexts. Within each subspace, we employed the Hill-Climbing (HC) greedy search algorithm for network structure learning. To evaluate the goodness of fit of continuous RNA-seq data, we used the Bayesian Gaussian Equivalence (BGe) score (Heckerman et al. 1995) as the objective function. The BGe score assumes a joint multivariate Gaussian distribution, allowing the calculation of the posterior probability of the network structure given the data. To capture temporal causal relationships, we imposed strict constraints on the network structure: only edges from time point t to time point t+1 (inter-time slice edges) were allowed, and intra-slice edges or reverse edges were prohibited. To assess the stability of edges, we further performed 25 Bootstrap resampling iterations (Efron and Tibshirani 1994) within each subspace. The final network retained only those regulatory relationships that were highly stable, defined as edges that were detected at a frequency of 1.0 (100% Consistency) in all related Bootstrap iterations.

## Data availability

All sequencing data generated in this study have been submitted to the NCBI Gene Expression Omnibus (GEO; https://www.ncbi.nlm.nih.gov/geo/) under accession numbers GSE3154440 and GSE316178. The sequencing data generated in this study have been submitted to the NCBI BioProject database (https://www.ncbi.nlm.nih.gov/bioproject/) under accession number PRJNA1394694.

## Code availability

All code to reproduce the analyses is available at: https://github.com/mingfengliu/DRGS.

## ACKNOWLEDGMENTS

We acknowledge the support of Theresa Nahreini and Emily Proksch at the Biochemistry Cell Culture Facility. We would like to thank Jes Persinger for their assistance with performing computational analysis on the Fiji cluster at the BioFrontiers Institute. T.R.C. is an investigator of the Howard Hughes Medical Institute.

## Author contributions

J. L. R. designed the study. M. L., S. S., I. T. P., A. R. S. P., and C. M. performed experiments. M. L., I. M., V. D., G. S., and A. W. performed bioinformatic analyses. J. L. R. provided access to resources. M. L. wrote the original draft. T. R. C. and J. L. R. supervised the study. All authors reviewed and approved the manuscript.

